# A two-dose regimen of Qβ virus-like particle-based vaccines elicit protective antibodies against heroin and fentanyl

**DOI:** 10.1101/2024.08.28.608988

**Authors:** Isabella G. Romano, Brandi Johnson-Weaver, Susan B. Core, Andzoa N. Jamus, Marcus Brackeen, Bruce Blough, Subhakar Dey, Yumei Huang, Herman Staats, William C. Wetsel, Bryce Chackerian, Kathryn M. Frietze

## Abstract

Opioid overdoses and the growing rate of opioid use disorder (OUD) are major public health concerns, particularly in the United States. Current treatment approaches for OUD have failed to slow the growth of the opioid crisis. Opioid vaccines have shown pre-clinical success in targeting multiple different opioid drugs. However, the need for many immunizations can limit their clinical implementation. In this study, we investigate the development of novel opioid vaccines by independently targeting fentanyl and the active metabolites of heroin using a bacteriophage virus-like particle (VLP) vaccine platform. We establish the successful conjugation of haptens to bacteriophage Qβ VLPs and demonstrate immunogenicity of Qβ-fentanyl, Qβ-morphine, and Qβ-6-acetylmorphine in animal models after one or two immunizations. We show that in independently or in combination, these vaccines elicit high-titer, high-avidity, and durable antibody responses. Moreover, we reveal their protective capacities against heroin or fentanyl challenge after two immunizations. Overall, these findings establish Qβ-VLP conjugated vaccines for heroin and fentanyl as very promising opioid vaccine candidates.

## INTRODUCTION

Opioid overdoses commonly involve prescription opioids, heroin, or fentanyl with each opioid drug class playing a major role in the rising overdose rates over the last several decades^1,2^. In the United States there were approximately 107,000 drug overdose deaths in 2022, with over 70% of those deaths attributed to opioids, totaling approximately 82,000 overdoses (National Institute on Drug Abuse, 2022). The majority of recent opioid deaths are associated with use of the potent synthetic opioid fentanyl either alone or in combination with other substances. While fentanyl has been a primary driver of the opioid crisis in recent years, nearly 20% of opioid overdose deaths involve heroin, emphasizing the continued relevance of both heroin and fentanyl in the opioid crisis (CDC 2022). Heroin, fentanyl and other opioids elicit analgesia, euphoria, anti-nociception, and respiratory depression through the activation of μ opioid receptors in the central nervous system^3,4^.

The most common interventions for opioid use disorder (OUD) and overdose management are medications for opioid use disorder (MOUD)^5^. These include opioid receptor antagonists such as naloxone and receptors agonists such as methadone and buprenorphine^6^. The stigma associated with MOUD, tight regulation, and the risk of physical dependence on the MOUD drugs pose challenges to this approach for management and limit patient accessiblity^7–11^. The large population of individuals with OUD experiencing housing insecurity or who lack of transportation services further exacerbates the challenges of patient accessibility to MOUD^12^. Naloxone (Narcan) is the current standard of care for reversal of opioid overdose. However, effectiveness of naloxone is dependent on bystander possession of the drug and administration in a time-sensitive manner^13,14^. Despite the approval and availability of MOUD, the opioid crisis continues to grow as a public health concern, emphasizing a need for the development of new treatments^15^.

The development of novel treatment options for OUD and overdose are of increasing interest as an area of focus including immunotherapies, such as monoclonal antibodies and vaccines^29,16^. Vaccines are a promising option because, unlike other treatments, they do not have abuse potential, may offer long-lasting protection, and do not hold the same negative stigma as MOUD. Vaccines against drugs of abuse have been investigated in pre-clinical and clinical studies for several targets including cocaine^18^, methamphetamine^19^, nicotine^20,21^ and heroin^22,23^. However, no vaccines for substance use disorders have obtained FDA approval for use in humans. Because drugs of abuse are poorly immunogenic on their own, vaccines classically involve linking a modified drug hapten to an immunogenic protein carrier [e.g., keyhole limpet hemocyanin (KLH) or tetanus toxoid (TT)], and formulation of the vaccine with a potent adjuvant^24,25^. An effective opioid vaccine requires the generation of durable, high-titer, high-affinity, serum antibodies that bind to and sequester opioid drug targets in the blood; thereby, limiting the amount of free drug that can cross the highly selective blood-brain barrier^26^. Previous studies have established the feasibility and pre-clinical efficacy of opioid-targeted vaccines to sequester these drugs in blood and offer protection from adverse effects of drug activity. After demonstrating an ability to elicit antibody responses and *in vivo* protective effects, some of these vaccine candidates have begun or are nearing clinical trials, including a KLH-based oxycodone vaccine^27–29^ (NCT04458545). Heroin-specific vaccines have been developed with a TT carrier protein conjugated to 6-monocacetylmorphine (6-AM) and formulated with Alum and CpG oligodeoxynucleotide adjuvants. These vaccines have shown efficacy in rodent and nonhuman primate models after three or more immunizations^30–33^. Fentanyl vaccine candidates have also been developed with the leading candidates using a cross-reactive material 197 (CRM-197) -based design and demonstrating promising pre-clinical protection with plans to enter clinical trials ^34–36^.

Virus-like particles (VLPs) are immunogenic, non-infectious, spontaneously self-assembling, highly repetitive and rigid structures formed by viral structural proteins that have been used as vaccine platforms or stand-alone vaccines to generate high-titer, durable antibody responses in few doses^37,38^. Qβ VLPs are derived from the icosahedral, RNA bacteriophage Qβ and have been used as a vaccine platform for a wide range of targets, including protein antigens from pathogens, self-antigens involved in chronic diseases, and small haptens (e.g., nicotine)^39–42^. Moreover, we previously have generated an oxycodone vaccine candidate using the Qβ VLP approach and demonstrated its immunogenicity and protective capacities in mouse models^43^. Qβ VLPs are thermostable and can be purified in high quantities with ease and low production costs. Its highly repetitive structure and particulate nature renders the Qβ VLP ideal for eliciting high-titer and durable antibody responses against small molecules and peptides in few immunizations^44,45^. Bacteriophage VLPs also possess several desirable features for clinical applications including compatibility with GMP manufacturing and an optimal safety profile in humans^46^. Here, we develop novel vaccines targeting heroin and fentanyl using a Qβ bacteriophage VLP design.

In this study, opioid targets were custom synthesized to allow for subsequent chemical conjugation onto the surface of Qβ VLPs to generate Qβ-morphine, Qβ-6-AM, and Qβ-fentanyl vaccines. We show these vaccines, independently and in combination, elicit high-titer, high-avidity, and durable antibodies in mice. Additionally, we show that two doses of Qβ VLP-based vaccines targeting heroin and fentanyl can blunt opioid-induced anti-nociception and respiratory depression in mice. Together, these data establish the feasibility of Qβ VLP-based vaccines against heroin and fentanyl as interventions for OUD and overdose.

## MATERIALS AND METHODS

### Chemical synthesis of morphine, 6-acetylmorphine and fentanyl haptens

Morphine-(Gly)_4_-Cys and 6-acetylmorphine-(Gly)_4_-Cys and Fentanyl-(Gly)_4_-Cys (FENBB3) hapten targets were custom synthesized at CellMosaic Inc. (Woburn, MA, USA). Morphine and 4ANPP (4-Aminophenyl-1-phenethylpiperidine) were purchased from Cayman Chemical Company Inc. (Ann Arbor, MI). Morphine was modified at CellMosaic to introduce a carboxylic acid functional group via the phenolic functionality. Carboxyl functional morphine is then acetylated to produce 6-acetyl-morphine (6-AM). 4ANPP (4-Aminophenyl-1-phenethylpiperidine) was converted to carboxy butyryl fentanyl via modification of the aromatic amino group. A custom Gly-Gly-Gly-Gly-Cys [(Gly)_4_-Cys)] peptide with protected Cys was synthesized using standard Fmoc Solid Phase Peptide Protocol. The *N*-terminal of the peptide contained a free amine group, and the C-terminal of the peptide was an amide. The peptide was coupled first to morphine acid, 6-acetylmorphine acid, or fentanyl acid based on similar literature protocols^47^. After removal of the Cys protecting group, the conjugate was purified by standard C18 HPLC using TFA/water/acetonitrile system and lyophilized to dryness. The identity of the conjugate was confirmed by MALDI-TOF or ESI MS. Calculated exact mass: 673.25 (morphine-(Gly)_4_-Cys). Obtained: [M+H]^+^: 674.5 (morphine-(Gly)_4_-Cys). Exact mass: 715.78 (6-acetylmorphine-(Gly)_4_-Cys). Obtained: [M+H]^+^: 716.4 (6-acetylmorphine-(Gly)_4_-Cys). Exact mass: 710.32 (Fentanyl-(Gly)_4_-Cys). Obtained: [M+H]^+^: 711.3 (Fentanyl-(Gly)_4_-Cys).

### Expression and purification of bacteriophage Qβ VLPs

Qβ bacteriophage VLPs were produced and purified as previously described^39–41^. Qβ VLPs were expressed from plasmid pETQCT using electrocompetent*. E.coli* C41 cells (#CMC0021; MilliporeSigma, Burlington, MA, USA). The 1:10 diluted plasmid was combined with C41 cells (∼3 x 10^9^ competent cells) and incubated on ice for 5 min followed by electroporation using a Bio-Rad Electroporator (Bio-Rad, Hercules, CA, USA). The transformation was incubated at 37°C in LB media for 1 hr prior to spreading onto LB agar plates containing kanamycin. A single colony was picked and incubated in LB broth with 50 μg/mL kanamycin at 37 °C until the OD_600_ was 0.6 followed by induction with 0.5 mM IPTG for 3 hr at 37°C. Bacterial pellets were obtained via centrifugation, re-suspended in lysis buffer (5.8 g NaCl, 3.7 g EDTA, 7.9 g Tris HCl in ddH_2_O) with 10% deoxycholate and incubated on ice for 30 min followed by sonication. Next, 10 mg/mL DNase and 2 M MgCl_2_ were added to digest the residual DNA. The supernatant was isolated by centrifugation and incubated in 60% ammonium sulfate overnight at 4 °C. Following centrifugation, the resulting pellets were re-suspended in cold Sepharose column buffer, ultra-centrifuged at 10,000 rpm, and the supernatant was fractionated by size-exclusion chromatography on a Sepharose column (Sepharose CL-4B; Global Life Sciences Solutions, Wilmington, DE, USA) to separate the Qβ containing fractions. Qβ containing fractions were incubated in 70% ammonium sulfate overnight at 4°C to precipitate the protein. Following ultracentrifugation, residual endotoxin was depleted using Triton X-114 and the final concentration of VLPs was determined via SDS-PAGE using known concentrations of hen’s egg lysozyme as a control.

### Conjugation of haptens to Qβ VLPs and preparation of vaccine doses

Hapten targets were independently conjugated to the surface of Qβ VLPs using the heterobifunctional cross-linker succinimidyl-6-([β-maleimidopropionamido]hexanoate) or SMPH (#22363, Thermo Fisher Scientific, Waltham, MA, USA). Qβ VLPs were incubated with SMPH at a molar ratio of 10:1 (SMPH: Qβ coat protein) for 2 hr at room temperature to permit the reaction with Qβ surface exposed lysine residues. Excess SMPH was removed by amicon filtration using an Amicon Ultra-4 centrifugal unit with a 100-kDa cutoff (#UFC810024, Amicon, Miami, FL, USA). Haptens were independently added to Qβ-SMPH at a molar ratio of 10:1 (hapten: Qβ coat protein) and incubated at 4°C overnight. Successful conjugation was confirmed via SDS-PAGE using a 12% denaturing gel (Invitrogen, Waltham, MA, USA) followed by Coomassie staining. Successful conjugation was determined as generating a shift upwards in molecular weight accompanied by a laddering pattern to indicate the attachment of multiple hapten targets per Qβ coat protein, with a reduction in unconjugated Qβ coat protein (14 kDa). Vaccine doses were diluted to a concentration of 20 μg/50 μL or 10 μg/50 μL for mouse immunizations. Combination vaccines were generated by combining Qβ-morphine + Qβ-6-AM or Qβ-morphine + Qβ-6-AM +Qβ-fentanyl using 20 μg or 10 μg of each hapten.

### Mouse immunization studies

All animal procedures were approved by the University of New Mexico (IACUC protocol #20-201045) and the Duke University Intuitional Animal Care and Use Committees (IACUC protocol A094-04-23). Male and female BALB/cJ mice (6-8 weeks old; Jackson Laboratories, Bar Harbor, ME) were used for immunizations. Animals received immunization (i.m.) via injection into the hind-legs. Animals receiving one immunization were vaccinated on day 0 and animals receiving a second immunization were vaccinated on days 0 and 21. Retro-orbital or submandibular bleeds were followed by centrifugation at 10,000 rpm in a microfuge for 10 min to harvest sera for subsequent analyses.

### ELISA to determine antibody titers

ELISAs to measure opioid-specific serum IgG antibodies were performed in a similar manner as reported by Jones and colleagues with minor modifications^48^. Specifically, 384-well Maxisorp ELISA plates (Thermo FisherScientific) were coated with the opioid-protein conjugate antigens at 2 μg/ml in carbonate/bicarbonate (CBC) buffer and incubated overnight at 4°C. ELISA plates were washed and blocked with CBC buffer containing nonfat dry milk for 2 hr at room temperature. Serially diluted (2-fold) serum samples were added to the ELISA plate at a starting dilution of 1:32 and incubated at 4°C overnight. The plates were washed before the addition of goat anti-mouse IgG secondary antibody conjugated with alkaline phosphatase (AP) (Southern Biotech, Birmingham, AL, USA) for 2 hr. ELISA plates were developed using the fluorescent ATTOPHOS substrate system (Promega, Madison WI, USA) for 15 min. A reference sample from naïve, unimmunized mice was included on each plate to calculate the endpoint titer. Here, the endpoint titer was defined as the last experimental sample dilution that provided a positive signal 3-fold greater than the naïve reference sample at the same dilution.

Ninety-six well microtiter plates (Immulon, #3455) were coated for 2 h at room temperature with 250 ng/well of morphine-BSA (#MBS537653; MyBioSource, San Diego, CA, USA), 6-AM BSA (#Vang-Cr3644; Creative Biolabs, Shirley, NY), or fentanyl-BSA (#DAG398; Creative Diagnostics, New York, NY, USA) in PBS followed by blocking with 0.5% milk/PBS solution (#A614-1005; Quality Biological, Gaithersburg, MD, USA) overnight at 4 °C. Serial dilutions of immune sera (4-fold increases from 1:40 to 1:655360) were added to each well and incubated at room temperature for 2 hr. HRP-conjugated goat anti-mouse IgG, 1:5000 in blocking solution (#15-035-003; Jackson ImmunoResearch, West Grove, PA, USA) was added and incubated for 45 min at room temperature. Following five washes, 50 μL of 3,3′,5,5′-tetramethylbenzidine (#EM613544; MilliporeSigma) (TMB) substrate was added and incubated for 10 min. 50 μL of 1% HCl (#AC12463-500 L; Thermo Scientific Chemicals, Waltham, MA, USA) terminated the reaction and absorbance was measured using a microplate reader at 450 nm (#AccuSkan FC; Thermo Fisher). Endpoint dilution IgG is reported as the final serum dilution that generated A450 greater than twice that of background.

### Surface Plasmon Resonance (SPR) measurements of serum antibody avidity

Sera avidity to antigens 6-AM BSA and morphine-BSA were quantitated by surface plasmon resonance (SPR) analysis (#BIAcore^TM^ 3000; BIAcore/GE Healthcare, Pittsburgh, PA, USA). The binding response was measured by SPR following immobilization of antigens^49^ on CM5 sensor chips (BIAcore/GE Healthcare). Mouse sera samples diluted 1:50 in 1x PBS, flowed (2.5 min) over the immobilized antigen surfaces followed by a dissociation phase (post-injection/buffer wash) of 10 min and a regeneration with glycine at pH 2.0. Non-specific binding of pooled negative control sera were subtracted from each of the post-immunization sera-sample binding-data. Data analyses were performed with BIA-evaluation 4.1 software (BIAcore/GE Healthcare). Binding responses were quantitated by averaging the post-injection response unit (RU) over a 10 sec window; the dissociation rate constant k_d_ (second^-1^), was determined during the post-injection phase after the stabilization of signal. A positive response was defined as when both replicates had a RU value ≥10. The relative avidity binding score was calculated as: Avidity score (RU.s) = (Binding Response Units /k_d_).^50^

### Anti-nociception behavioral studies

The anti-nociceptive effects of heroin or fentanyl (NIDA Drug Distribution Program, Bethesda, MD, USA) were assessed using thermal sensitivity in a tail-flick (Columbus Instruments) assay^51^. The laser on the tail flick apparatus was set to an intensity of 11, with a cut-off exposure time of 20 sec. Baseline responses were collected for tail-flick prior to opioid exposure. Vaccine-induced inhibition of the anti-nociceptive effects of heroin or fentanyl were monitored in immunized mice 3 weeks after the last immunization for each vaccine condition. Heroin (0.5 mg/kg, s.c.) or fentanyl (0.0625 mg/kg, s.c.) were administered. At 15, 30-, 60-, 90-, and 120-min post-injection, assessments of tail flick responses were conducted. The latencies for the mouse to flick its tail were quantitated. The data are expressed as the percent maximal possible effect (% MPE) for tail-flick as [(drug – basal response)/(20 sec – basal response time)] x 100%.

### Whole body plethysmography-based study of respiratory depression

Fentanyl-induced respiratory depression was assessed in vaccinated mice (n=8, BALB/cJ, male/female) 3-weeks post-second immunization. In the week leading up to drug challenge, mice were acclimated to respiratory chambers of the whole-body plethysmography (WBP) system (Buxco Data Sciences International, St Paul, MN, USA) daily for 30-min. On the day of fentanyl challenge, baseline respiratory parameters were collected including respiratory frequency (breaths per min), tidal volume (mL of air), and minute volume (frequency * tidal volume; mL/min) over a 30-min baseline acclimation period. Once steady-state respiration was established, fentanyl citrate (0.25 mg/kg, s.c.) was administered (#NDC 63323-806-12; Fresensius Kabi USA, LLC, Wilson, NC). Thirty min later, animals were placed into the respiratory chambers and respiratory data were collected over a 90 min time-course. Data collection was conducted using a WBP AHR universal study with pull bias flow mode. Measurements were recorded every 2 sec and expressed as a 60-sec rolling average. Data are shown with male and female animals combined for vaccinated and control mice as well as animals separated by sex in each vaccine condition.

### Statistics

Vaccine-induced serum antibody responses and nociception assays were compared using the nonparametric Mann-Whitney and Kruskal-Wallis tests with multiple comparisons to PBS or Qβ -immunized animals. Respiratory data were analyzed using a Two-way ANOVA mixed effects analysis with Sidak’s multiple comparison test. All statistical tests were performed using GraphPad Prism. A p < 0.05 was considered statistically significant for all results.

## RESULTS

### Generation of haptens and conjugation to Qβ VLPs

Qβ VLPs are made up of 180 copies of coat protein with 4 surface-exposed lysine residues on each copy, totaling 720 available positions for conjugation (**Figure 1A)**^52,53^. Conjugation of hapten targets is accomplished via the bifunctional chemical cross linker SMPH that reacts with surface-exposed lysine residues of the VLP on one arm and on the opposite arm to the free reactive sulfhydryl group of terminal cysteine residues on hapten targets **(Figure 1A)**. To design fentanyl targets for conjugation, three different linker compositions were tested: polyethylene glycol (PEG; FENBB1), a straight carbon chain ((CH_2_)_n_ ;FENBB2), and peptide ((Gly)_4_-Cys; FENBB3). All three linkers contained a terminal cystine to permit chemical conjugation to SMPH. These three linkers **(Figure 1B)** were examined to identify the optimal target for fentanyl vaccine design and to test other linkers as alternatives to the peptide linker used in a previous VLP-based opioid-vaccine design^43^. All three fentanyl targets conjugated successfully to the Qβ coat protein, but with varying efficiencies. The PEG and CH_2_ linkers (FENBB1 and FENBB2, respectively) resulted in only a single copy of hapten attached per coat protein with a significant amount of unconjugated coat protein (14 kDa band) remaining. The peptide linker ((Gly)_4_ Cys), FENBB3) showed the highest degree of conjugation with multiple copies of drug per individual coat protein with little-to-no unconjugated Qβ coat protein remaining.

**Figure 1.**
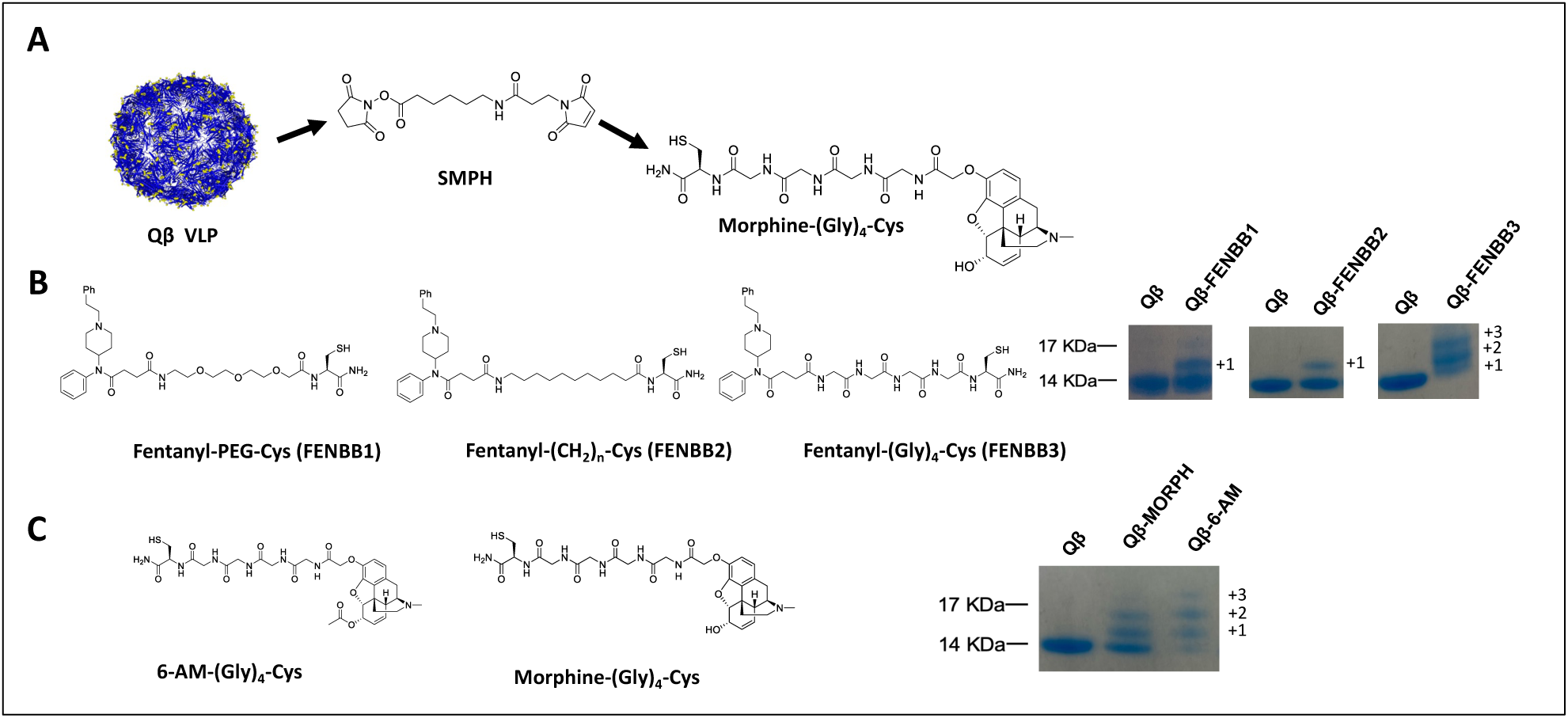
Hapten target design and conjugation to Qβ coat protein. **(A)** Hapten targets are chemically conjugated to Qβ VLPs using the bifunctional chemical crosslinker SMPH. Yellow dots indicate surface-exposed lysine residues on Qβ VLPs available for conjugation. Morphine-(Gly)_4_ Cys is shown as a representative example; all haptens are conjugated in the same manner. **(B)** Hapten design of 6-acetylmorphine (6-AM) and morphine-(Gly)_4_ Cys targets. Successful conjugation shown by Coomassie stained SDS-PAGE. Unconjugated Qβ coat protein has a MW of 14 kDa. The banding pattern of migration indicates 1, 2, or 3 copies of hapten attached per coat protein. **(C)** Fentanyl hapten targets using various linkers for attachment. Coomassie stained SDS-PAGE shows successful conjugation as in **(B).**

In the body, heroin is rapidly metabolized in two sequential deacetylation reactions to produce the active metabolites 6-monoacetylmoprhine (6-AM) and morphine^54,55^. It was believed previously that morphine was the primary metabolite responsible for the neural activity of heroin^56,57^. However, recent studies have shown that 6-AM is primarily responsible for the rapid neural effects of heroin^58,59^. Antibodies to these metabolites are effective in blocking heroin challenge and therefore were targeted in the development of our VLP based vaccine against heroin^30,31,60^ **(Figure 1C)**. In our heroin vaccine design, we utilized a peptide based (Gly)_4_ Cys linker that showed the highest degree of conjugation among the different linkers tested among our fentanyl vaccine candidates (**Figure 1B**). Moreover, this linker had been used by our group and others to display peptide and drug targets on VLPs or other carrier proteins^24,30,41,60–62^. Successful conjugation to Qβ VLPs was confirmed by SDS-PAGE with Coomassie staining. An increase in the apparent molecular weight and a laddering pattern compared to unconjugated Qβ coat protein (**Figure 1C**) confirmed the successful generation of Qβ VLPs displaying an estimated 120-170 copies of hapten per VLP.

### Qβ-morphine and Qβ-6-AM generate high-titer, high-avidity, and durable antibody responses in mice

Mice were immunized on day 0 and day 21 with Qβ-morphine, Qβ-6-AM, or Qβ-Combo (Qβ-morphine + Qβ-6-AM) vaccine candidates along with PBS and unconjugated Qβ controls. Male and female mice immunized with Qβ-morphine, Qβ-6-AM, or Qβ-Combo generated ∼10^7^ anti-morphine IgG endpoint titers (**Figure 2A)** and ∼ 10^4^ anti-6-€acetylmorphine IgG endpoint titer (**Figure 2B)**. We hypothesized that immunization would generate high-avidity antibody responses, consistent with classical prime-boost vaccine strategies^40,63^. To study antibody affinity, we used surface plasmon resonance (SPR) with immune sera collected at day 28 post-first immunization (day 7 post-second). Serum avidity is expressed as binding response units/K_D._ Avidity was determined to both morphine-BSA and 6-AM-BSA conjugate antigens in each vaccine group. All vaccines groups generated high avidity antibodies to morphine and 6-AM targets compared to control groups **(Figure 2C).** Across each vaccine group, antibody avidity to morphine was consistently higher than avidity to 6-AM.

**Figure 2.**
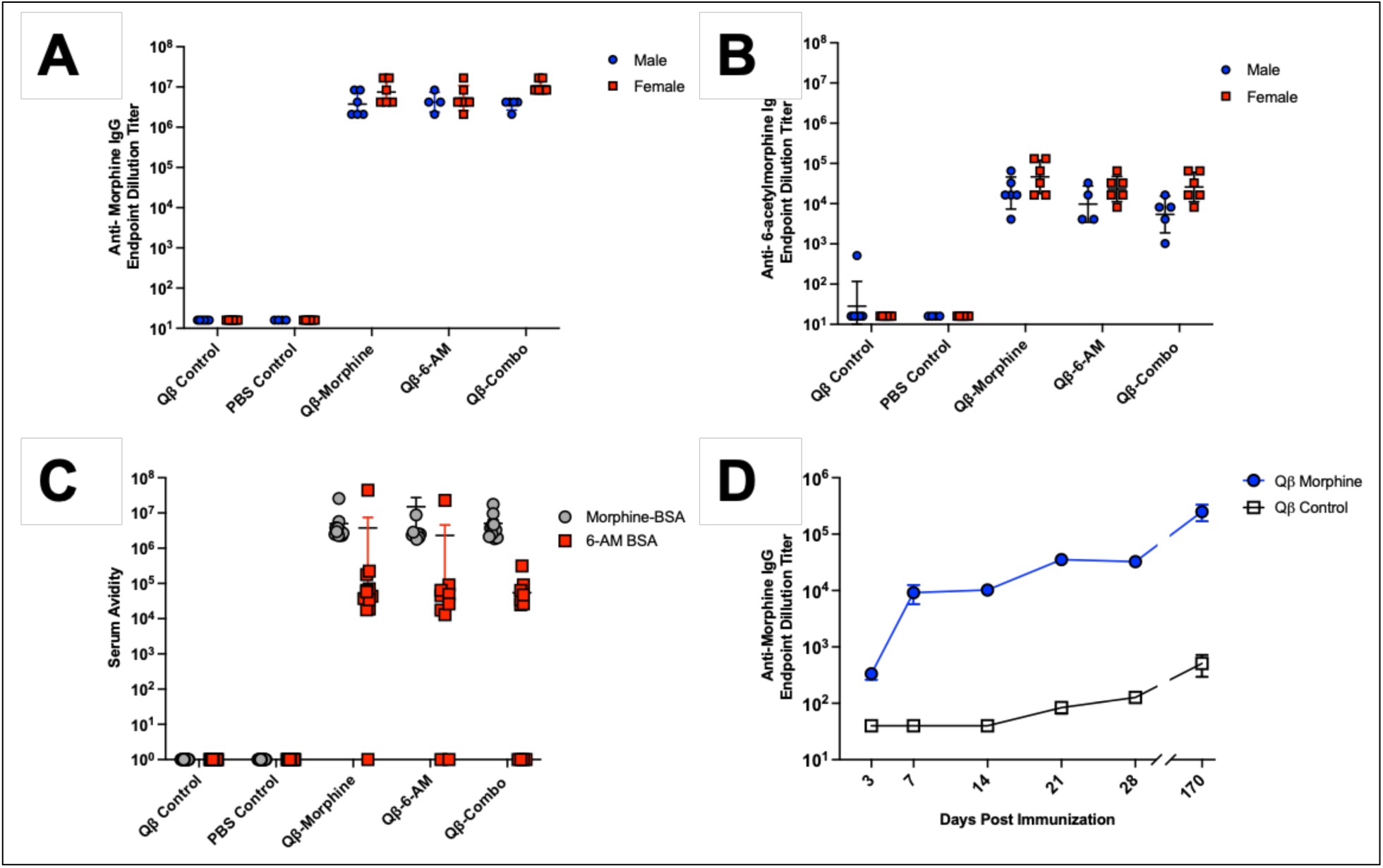
Qβ-morphine and Qβ-6-acetylmorphine vaccine candidates generate high titer, high avidity, durable antibody responses. Endpoint dilution IgG titers of vaccine-elicited anti-morphine **(A)** and anti-6-acetylmorphine **(B)** antibody responses after two immunizations. Responses generated by immunization with Qβ-morphine, Qβ-6-AM, or Qβ-Combo at Day 28 (Day 7 post-second immunization) are indicated. Data are expressed as geometric mean titer and SD. No significant differences were observed between sexes; Mann-Whitney test. **(C)** Antibody avidity to morphine-BSA or 6-AM BSA targets measured by surface plasmon resonance. Serum Avidity is expressed as binding response units/ K_D._ **(D)** Endpoint dilution IgG titers of vaccine-elicited anti-morphine antibody responses after a single immunization (Day 0) with Qβ-morphine and monitored over 170 days; p=0.0043, Mann-Whitney tesr

To investigate the kinetics and durability of the generated antibody responses, a time-course study was conducted in mice given a single dose of Qβ-morphine. Immunization rapidly elicited a >10^4^ end-point dilution antibody titer against the cognate drug target and continued to increase over time (**Figure 2D)**. Animals were followed for 170 days and antibody titers did not decrease, establishing that antibody responses are durable after a single intramuscular immunization in mice without exogenous adjuvant.

### Fentanyl vaccines generate high-titer antibodies

Mice were immunized on days 0 and 21 with each fentanyl vaccine candidate independently. Sera were collected on days 3, 7, 21 28, 35 and 42 post-first immunization for each candidate along with animals vaccinated with the unconjugated Qβ control. Each fentanyl vaccine generated high-titer antibody responses, with an endpoint dilution IgG titers of ∼10^4^ at day 42 (**Figure 3A**). Because we did not observe differences in the immunogenicity of the fentanyl vaccines, we focused on FENBB3 for remaining experiments due to its increased conjugation efficiency compared to other targets.

**Figure 3.**
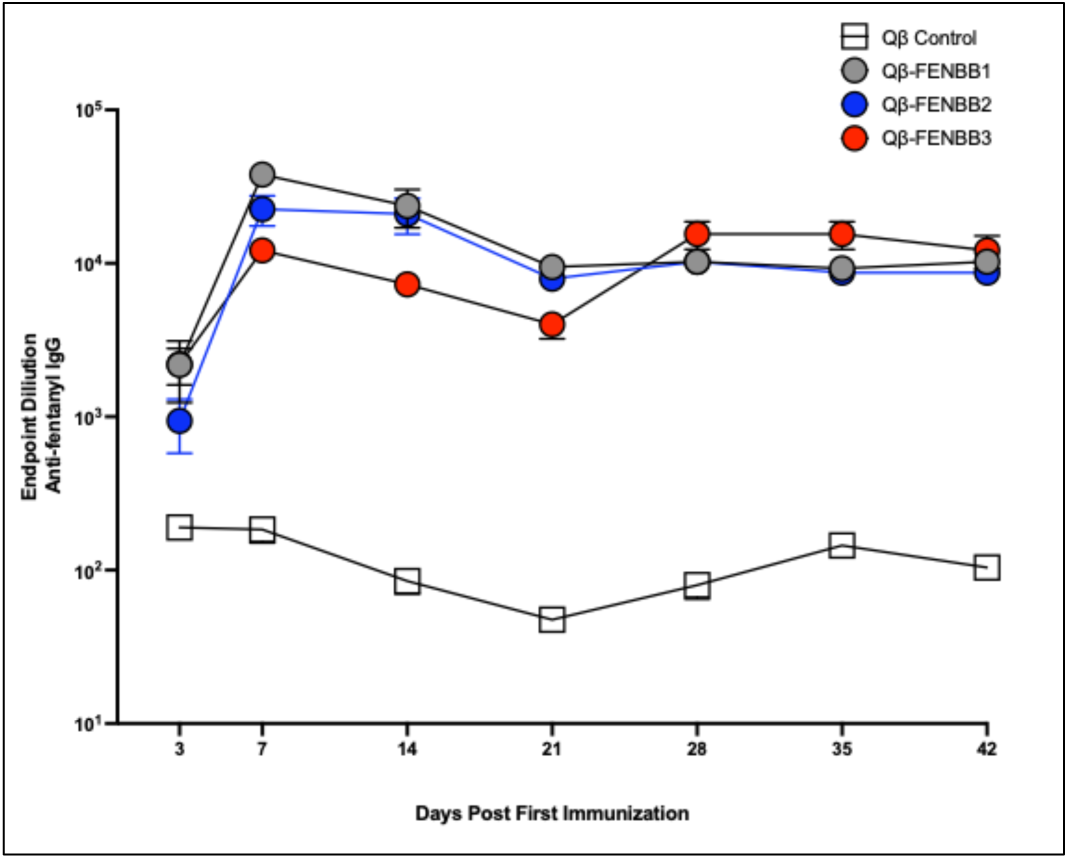
Qβ-fentanyl vaccines generate high-titer antibodies. Anti-fentanyl endpoint dilution IgG titers generated after immunization with FENBB1, FENBB2 or FENBB3 fentanyl vaccine candidates or with the unconjugated Qβ control. Mice (n=10, Balb/cJ, male and female) were immunized on days 0 and 21.

### Immunization with heroin vaccines offer significant protection from heroin-induced anti-nociception

Opioids generate anti-nociceptive effects, blocking pain signal transmission and subsequent physiological responses. The tail-flick anti-nociception assay is a common preclinical behavioral test to assess pain responses in animals and was used here to determine if Qβ vaccine candidates can blunt heroin-induced anti-nociception upon *in vivo* challenge. Based on a previous heroin dose-response study (data not shown), vaccinated animals were challenged with heroin (0.5 mg/kg, s.c.) at 3-weeks post-second immunization. Pain responses were tested via tail flick every 15 min for the first 30 min and followed at 30 min intervals to 90-min post-injection. Results are expressed as percent maximum possible effect (% MPE) calculated as the relationship of the latency to respond (withdrawal of tail from heat source) compared to baseline responses without drug. We show that vaccinated animals displayed attenuated heroin-induced anti-nociception across the entire time-course (**Figure 4A**). At 30-min post-injection, vaccinated animals showed significant protection from heroin-induced anti-nociception compared to the PBS control. Qβ-6-AM (p=0.0364) and Qβ-combo (p=0.0003); Dunn’s multiple comparison test (**Figure 4B**). Conversely, a single immunization did not provide sufficient protection from heroin challenge (**Supplemental Figure 1**). This demonstrates the requirement for two vaccine doses to achieve protective effects against cognate drug challenge.

**Figure 4.**
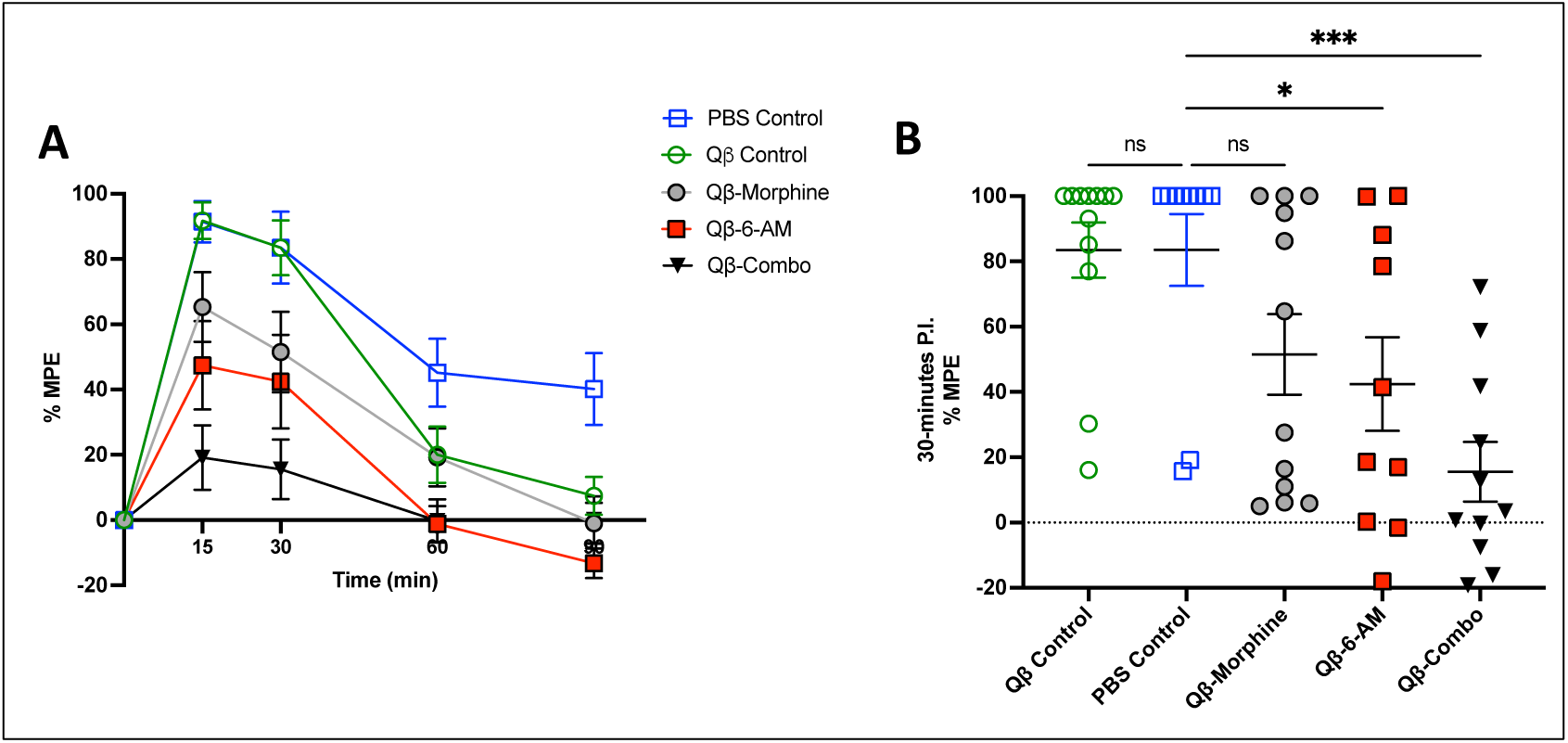
Qβ vaccines against heroin metabolites protect against heroin-induced anti-nociception. **(A)** Anti-nociceptive responses measured by tail-flick assay in Balb/cJ mice immunized with Qβ-morphine, Qβ-6AM, Qβ-Combo, Qβ control, or PBS control challenged with heroin (0.5 mg/kg, s.c.) at 3-weeks post-second immunization and tested over a 90-min time course. % Maximum Possible Effect (%MPE=[(drug – basal response)/(20 sec – basal response time)] x 100%). (B) Tail-flick anti-nociceptive responses analyzed at 30-min post-injection with heroin (0.5 mg/kg, s.c.). *p=0.0364; ***p=0.0003; *vs*. PBS control; Kruskal-Wallis test, GraphPad Prism.

### Qβ-fentanyl vaccines show protection from fentanyl-induced anti-nociception and respiratory depression

Mice immunized with Qβ-fentanyl (FENBB2; CH_2_ linker) were challenged with fentanyl (0.0625 mg/kg, s.c.) at 3-weeks post-second immunization. Parenthetically, the fentanyl dose was determined by a previously conducted pilot dose-response study (data not shown). Animals were assessed by tail-flick assay for anti-nociceptive responses starting 15-min post-injection over a 90-min period. Anti-nociceptive responses are reported as % MPE, as described for the heroin tail-flick assay above. Here, we show that over the observed time-course, Qβ-fentanyl vaccinated mice display significantly lower fentanyl-induced anti-nociception compared to unconjugated Qβ and PBS control mice at 15, 30 and 60-min post-injection; p<0.05, Tukey’s multiple comparison (**Figure 5A**)

**Figure 5.**
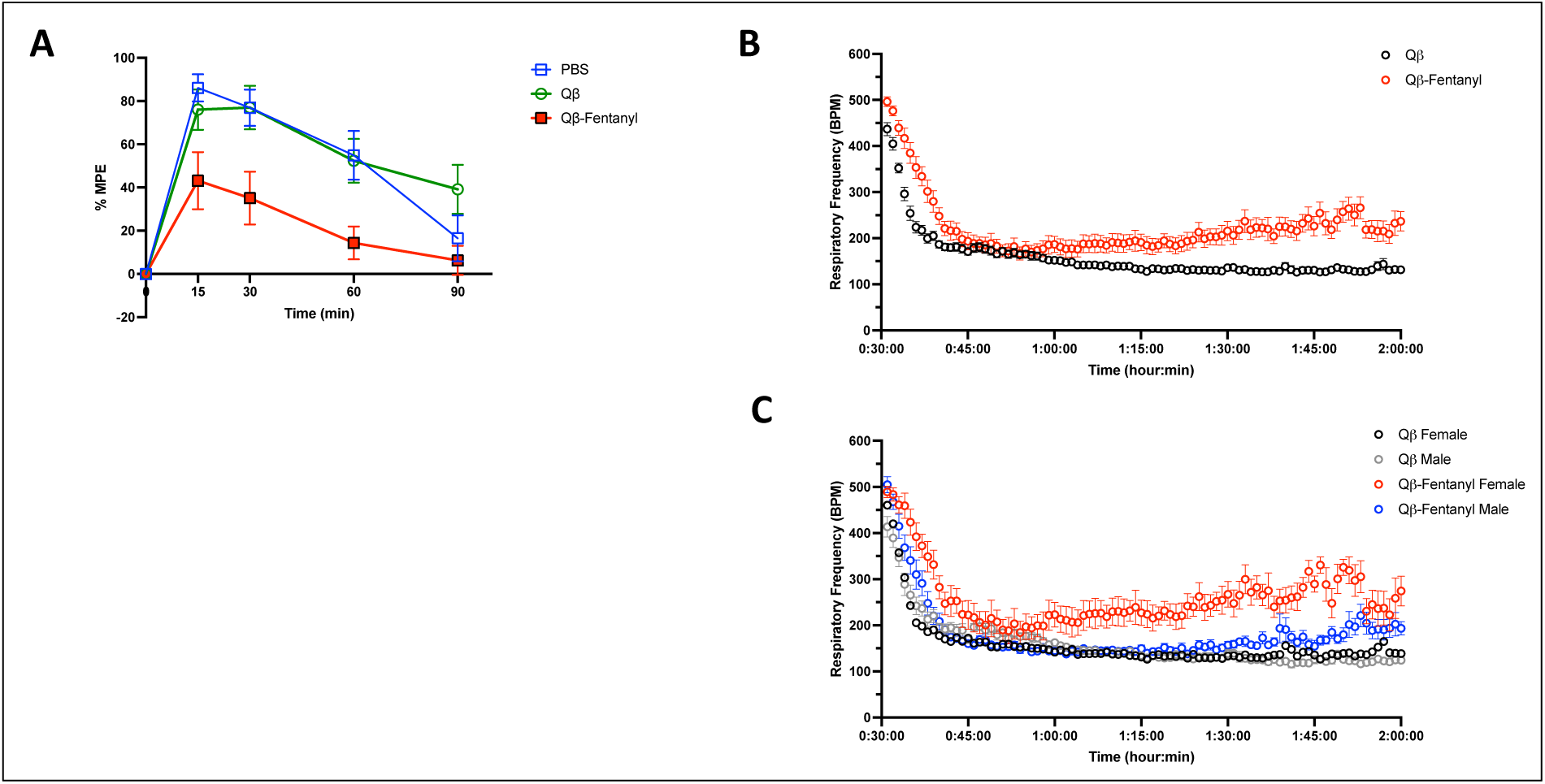
Qβ-fentanyl vaccines protect against anti-nociception and respiratory depression in mice. **(A)** Anti-nociceptive responses assessed by the tail-flick assay in mice vaccinated with Qβ-fentanyl, Qβ control, or PBS and challenged with fentanyl (0.0625 mg/kg, s.c.) at 3 weeks post-second immunization. % Maximum Possible Effect (%MPE=[(drug – basal response)/(20 sec – basal response time)] x 100%). P<0.05 at t= 15, 30, 60 and 90 compared to Qβ control. p<0.05 at t=15, 30 and 60 min compared to PBS control; Tukey’s multiple comparison. **(B/C)** Fentanyl-induced decline in respiratory frequency (breaths per minute; BPM) measured by whole-body plethysmography in vaccinated mice (3-weeks post-second immunization) challenged with fentanyl (0.25 mg/kg, s.c.) at 30 min after injection and recorded over 2 hr. Male and female mice (Balb/cJ; n=8) grouped together; p=0.0002; Mixed-Effects Analysis; GraphPad Prism **(B)** and separated by sex **(C)**.

Opioids cause respiratory depression which can lead to fatal overdose due to drug activity in the respiratory centers of the brainstem^64,65^. Fentanyl causes this phenomenon at lower doses than other opioids, making respiratory depression a particularly relevant endpoint for assessing vaccine-mediated protection against this opioid. To determine protection from fentanyl-induced respiratory depression, Qβ-fentanyl vaccinated mice were challenged with fentanyl (0.25 mg/kg, s.c.) at 3 weeks post-second immunization. The dose was determined based upon previously conducted pilot dose-response studies (data not shown). Animals were first acclimated to respiratory chambers in the whole-body plethysmography system for 30 min prior to drug administration. Following administration, animals were monitored over 90 min post-injection. We observed a lower magnitude of decrease in respiratory frequency accompanied by a faster time to recovery in mice vaccinated with Qβ-fentanyl compared to Qβ control (male and female mice shown together, **Figure 5B**). Notably, there was a clear sex-difference in protection observed as shown in **Figure 5C**, where overall protection was driven primarily by the responses of female mice (p=0.0002, Mixed-Effects Analysis), with male mice beginning to show recovery later than female mice but sooner than control mice.

### Combining heroin and fentanyl vaccines does not diminish morphine or fentanyl antibody responses but may introduce cross-reactivity to off-target opioids

Due to the increasing prevalence of polysubstance abuse and the concern for fentanyl contamination in the heroin drug circulation, we investigated the feasibility of combining heroin and fentanyl vaccine candidates to generate a mixed vaccine. To accomplish this, Qβ-morphine, Qβ-6AM, and Qβ-fentanyl (FENBB3; peptide linker) were combined and delivered in a single formulation. Vaccines were combined at a high dose of 20 μg or a lower dose of 10 μg of each of the three candidates and administered via injection (i.m.) to male and female BALB/cJ mice. Animals were immunized on day 0 and day 21, followed by sera collection and ELISA on day 28 (Day 7 post-second immunization). ELISAs were conducted using Fentanyl-BSA (**Figure 6A),** morphine-BSA **(Figure 6B),** naltrexone-BSA **(Figure 6C),** methadone-BSA **(Figure 6D),** or buprenorphine-BSA **(Figure 6E)** as coating antigens. Administration of the trivalent vaccine at 20 μg or 10 μg doses does not diminish antibody responses to fentanyl (**Figure 6A)** or morphine (**Figure 6B)** target antigens compared to fentanyl or heroin vaccines alone. Notably, heroin vaccines alone displayed minimal response to fentanyl, while fentanyl vaccines alone showed some cross-reactive antibody responses to morphine. The addition of Qβ-morphine and Qβ-6AM increased the cross-reactive responses to naltrexone and buprenorphine compared to Qβ-fentanyl alone (**Figure 6C, E**). Mild cross-reactivity was observed to methadone (**Figure 6D)** across all vaccine conditions. Importantly, off-target responses were several orders of magnitude lower than on-target responses for each vaccine condition tested.

**Figure 6.**
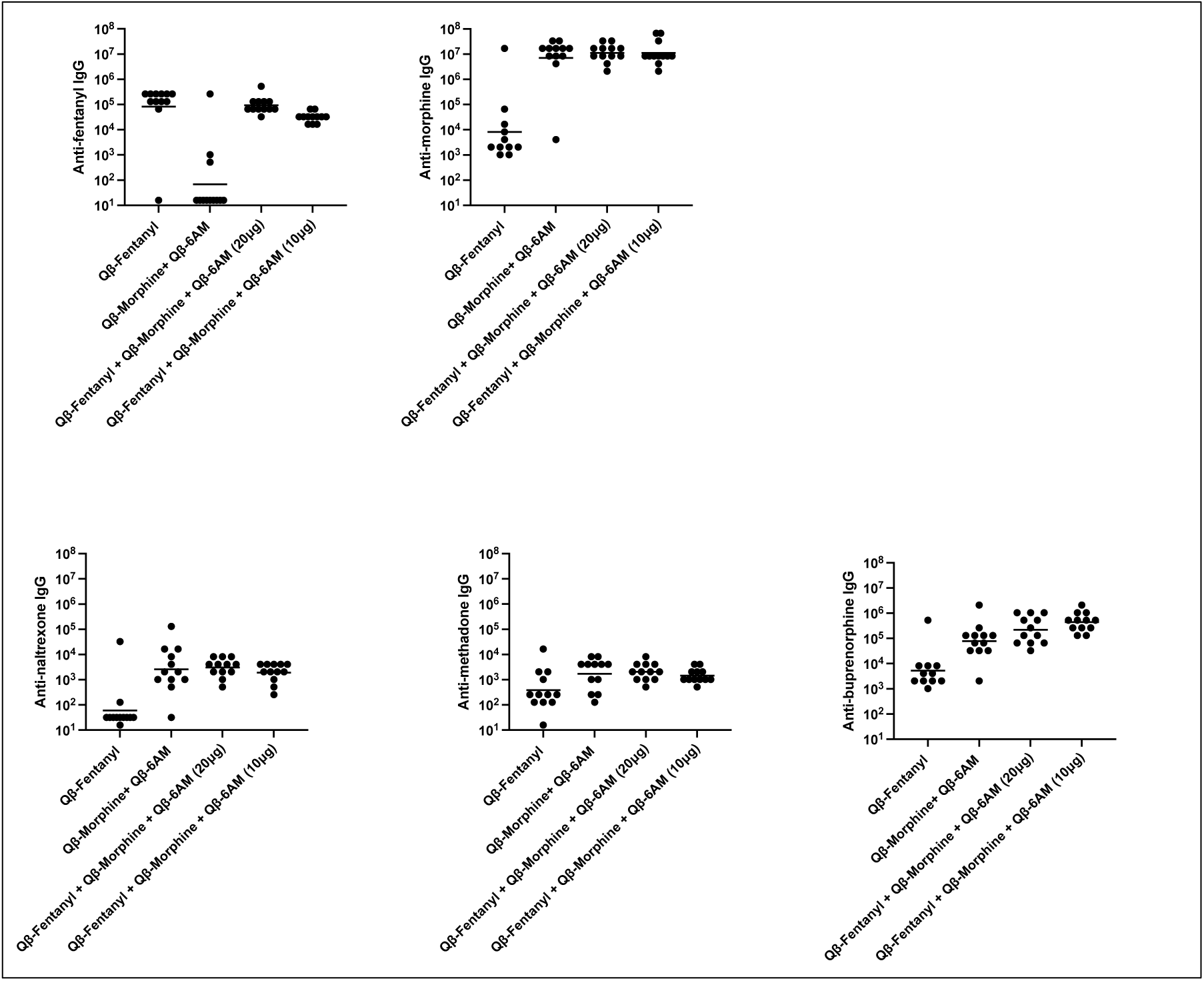
Combined fentanyl and heroin vaccine formulation elicits high-titer antibody responses with some cross-reactivity. Endpoint dilution IgG titers (geometric mean titer) against fentanyl **(A)**, morphine **(B)**, naltrexone **(C)**, methadone **(D)**, and buprenorphine **(E).** Mice (Balb/cJ, n=12, male and female) were immunized twice with Qβ-fentanyl, Qβ-morphine + Qβ-6-AM, or Qβ-fentanyl + Qβ-morphine + Qβ-6-AM (20 μg or 10 μg doses). Antibody titers were assessed at Day 28 (Day 7 post-second immunization).

### Combining heroin and fentanyl vaccines in a single formulation does not diminish protection from fentanyl-induced anti-nociception

We were interested in examining the impact of the addition of Qβ-based heroin vaccines into the previously established Qβ-fentanyl vaccine on vaccine-mediated protection from fentanyl-induced anti-nociception. Mice were challenged with fentanyl (0.0625 mg/kg, s.c.) at 3 weeks post-second immunization with Qβ-fentanyl, Qβ-morphine + Qβ-6-AM, Qβ-fentanyl + Qβ-morphine + Qβ-6-AM (20 μg or 10 μg doses), or PBS control. The addition of Qβ-morphine + Qβ-6-AM at high or low doses did not adversely affect the protection generated by Qβ-fentanyl against fentanyl-induced anti-nociception (**Figure 7**). All vaccine groups other than the heroin vaccine alone significantly reduced the impact of fentanyl, demonstrating the feasibility of combining multiple vaccine targets in a single immunization.

**Figure 7.**
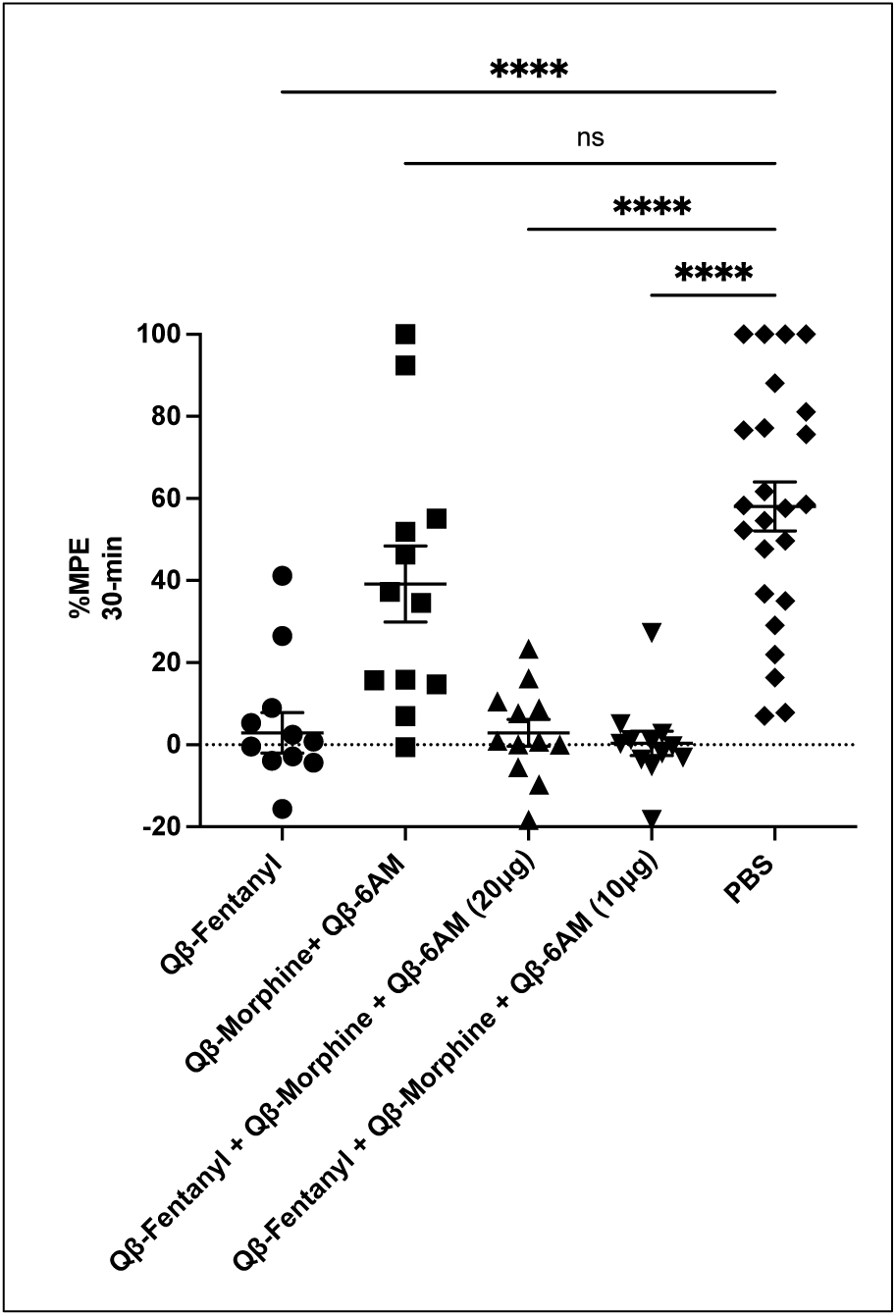
Combined fentanyl and heroin vaccine formulation shows protection against fentanyl-induced anti-nociception. Fentanyl-induced anti-nociception measured by tail-flick assay in vaccinated mice challenged with fentanyl (0.0625 mg/kg, s.c.) at 3 weeks post-second immunization. % Maximum Possible Effect (%MPE=[(drug – basal response)/(20 sec – basal response time)] x 100%). Responses at 30 min post-injection are shown. ****p<0.0001; Dunn’s multiple comparison test; GraphPad Prism.

## DISCUSSION

The growing opioid crisis has led to an increased interest in the development of novel treatments for OUD and opioid overdose prevention. Recently, opioid vaccines have been investigated, including a Qβ VLP-based vaccine for oxycodone as developed by our group ^26,27,29,43,66^. A related approach was published recently, using mutant Qβ to develop a heroin vaccine, which further provided feasibility for virus-like particle-based vaccines targeted against opioids^67^. In the present study, we report our efforts to develop Qβ VLP-based vaccines for heroin and fentanyl. Heroin vaccines were generated through the addition of a (Gly)_4_-Cys peptide linker and subsequent conjugation of hapten targets onto Qβ VLPs. We found the generated Qβ-morphine and Qβ-6AM vaccine candidates induced high-titer, high-avidity serum antibodies in mice when injected alone or in combination. The Qβ-morphine vaccinated mice were followed for a total of 170 days after a single administration and antibody titers remained high, demonstrating the durability of responses. Moreover, we demonstrated also the protective capabilities of these vaccine candidates to block heroin-induced anti-nociception upon *in vivo* drug challenge.

The fentanyl vaccine candidates were generated in a similar manner with the additional investigation of three different potential linker compositions for hapten target generation: polyethylene glycol (PEG; FENBB1), a straight carbon chain ((CH_2_)_n_ ;FENBB2), and peptide ((Gly)_4_ Cys; FENBB3). Alternate linker compositions were tested to determine if linkers other than the peptide(Gly)_4_ Cys utilized in heroin and oxycodone vaccine designs could be used to improve conjugation, immunogenicity, or protection^43^. We showed the successful generation and conjugation of each fentanyl target and the subsequent immunogenicity in mouse models. Two of the targets--FENBB2 and FENBB3--were selected for further assessment of vaccine-mediated protection and cross-reactivity. These candidates were selected primarily due to their higher conjugation efficiency and increased solubility compared to the PEG linker-based candidate. In further tests, we found animals vaccinated with Qβ-FENBB2 had protection from opioid-induced anti-nociception and demonstrate the high specificity of the vaccine-elicited antibodies with no significant cross-reactivity to methadone, buprenorphine or naltrexone. Moreover, animals vaccinated with Qβ-FENBB3 showed protection from fentanyl-induced respiratory depression, thereby demonstrating the protective capacities of the Qβ VLP-based vaccines targeting fentanyl.

The increasing prevalence of drug combinations, particularly the addition of fentanyl into the heroin supply, is a major concern for the rising rates of overdose and deaths. These dire circumstances are pleas for the development of effective new treatments^1^. To address this condition, we combined our heroin and fentanyl vaccine targets and determined that the combination of these two candidates did not reduce vaccine-elicited antibody responses or protection from fentanyl challenge. However, the addition of the heroin vaccine targets increased cross-reactive antibody responses to buprenorphine, methadone, and naltrexone. Cross-reactivity is an important issue when considering the development of opioid vaccines, with concerns regarding vaccinated patients’ ability to respond to the MOUD standard of care treatments for OUD. Other studies have established that vaccine-elicited antibodies do not block *in vivo* functions of MOUD^68,69^. In the present study, fentanyl vaccines alone did not generate significant cross-reactive antibodies; however, the addition of morphine and 6-AM vaccine targets appeared to drive cross-reactivity. Based upon the relatively low titer of cross-reactive antibodies compared to on-target antibodies, it is unlikely that this *in vitro* result will lead to significant *in vivo* cross-reactivity. Nevertheless, it will be important to investigate *in vivo* cross-reactivity in the future to determine the translational feasibility of combination vaccines and to demonstrate that vaccine-elicited antibodies will not limit the efficacy of MOUD.

Published preclinical studies involving opioid vaccine candidates have largely neglected to incorporate both male and female animals, with most of the previous research focused on males. In humans, men have a higher incidence of OUD and opioid overdose^70^. However, there are key sex differences that impact opioid activity and mediate sex differences in the effects of opioid receptor activation^71–74^. With the goal of developing clinically relevant heroin and fentanyl vaccines, it is imperative that both sexes be included in pre-clinical studies. Due to this past limitation, we included male and female mice in all aspects of our investigations. We found some minor differences in antibody responses and protection from anti-nociception; however, all tested vaccine candidates were immunogenic and protective from anti-nociception at low doses in both sexes (**Supplemental Figures 2 and 3**). At higher doses of a fentanyl challenge (0.25 mg/kg, s.c.), we observed a sex-bias in protection of respiratory depression, with female mice driving the effect. This result is intriguing, and it will require continued investigations in the future, especially when considering factors such as estrous cycle phase and opioid metabolism that could underlie the sex-specific differences we observed in our studies. Overall, it is critically important to include male and female animals in opioid vaccine efforts to identify possible sex differences in preclinical studies that will inform clinical trials.

In this study, we selected not to use exogenous adjuvants in our vaccine design. Qβ VLPs offered a more advantageous approach by its endogenous adjuvating activity mediated by the coding RNA encapsulated within the VLP acting as a toll-like receptor ligand to boost T helper responses and increase immunogenicity^37^. However, exogenous adjuvants (e.g., Advax and mastoparan-7) have been established to increase immune responses to both VLP- and non-VLP based vaccines for drugs of abuse^40,75^. Recently, A TLR7/8 agonist was shown to significantly improve the vaccine efficacy of anti-fentanyl vaccines^76^. It has been shown also that while IgG is the predominate isotype in blood and should play a significant role in drug sequestration within blood, IgA also plays an important role in mediating opioid vaccine protection^35^. To improve or skew antibody responses and therefore protection in our heroin and fentanyl vaccine designs, we may utilize adjuvants or alternate routes of immunization in the future.

In conclusion, we establish Qβ VLP based vaccines targeting heroin and fentanyl as novel opioid vaccine candidates.

## ACKNOWLEDGEMENTS

We thank Mr. Christopher Means and Ms. Julia Hoyt in the Mouse Behavioral and Neuroendocrine Analysis Facility for conducting the tail flick assays at Duke University.

## FUNDING

Research reported in this publication was supported by *Helping to End Addiction Long-term*^®^ *Initiative, or NIH HEAL Initiative*^®^ under award number UG3DA053123, NIH/NIDA F31DA059236, NIH/NIGMS P20GM109089, NIH/NCATS UL1TR001449, and NIH/NCATS KL2TR001448. This work was supported in part by the Dedicated Health Research Funds from the University of New Mexico School of Medicine.

## Notes

### Competing Interest Statement

B. Chackerian has equity in Metaphore Biotechnologies. All other authors have no competing interests.

