## Supplemental Figures for "A two-dose regimen of Qβ virus-like particle-based vaccines elicit protective antibodies against heroin and fentanyl"

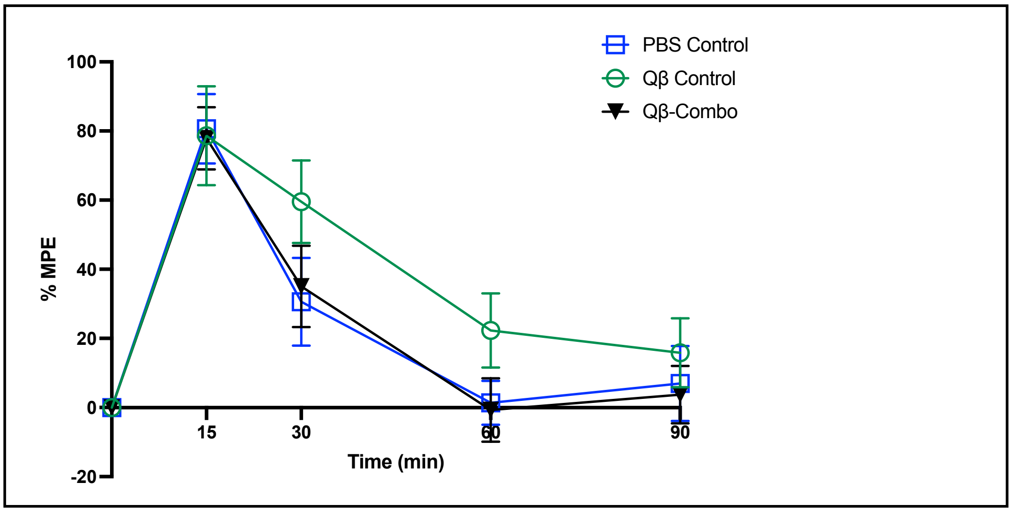


**Supplemental Figure 1. A single vaccine dose does not protect against heroin challenge.** Anti-nociceptive responses measured by tail flick in vaccinated mice (n=12, BALBc/j) challenged with s.c. heroin (0.5mg/kg) at three-weeks post-first immunization. Qβ-Combo= Qβ-Morphine + Qβ-6-AM. Data shown as mean and SEM. % Maximum Possible Effect (%MPE=[(drug – basal response)/(20 sec – basal response time)] x 100%). NS; Dunn's multiple comparison test; GraphPad Prism.

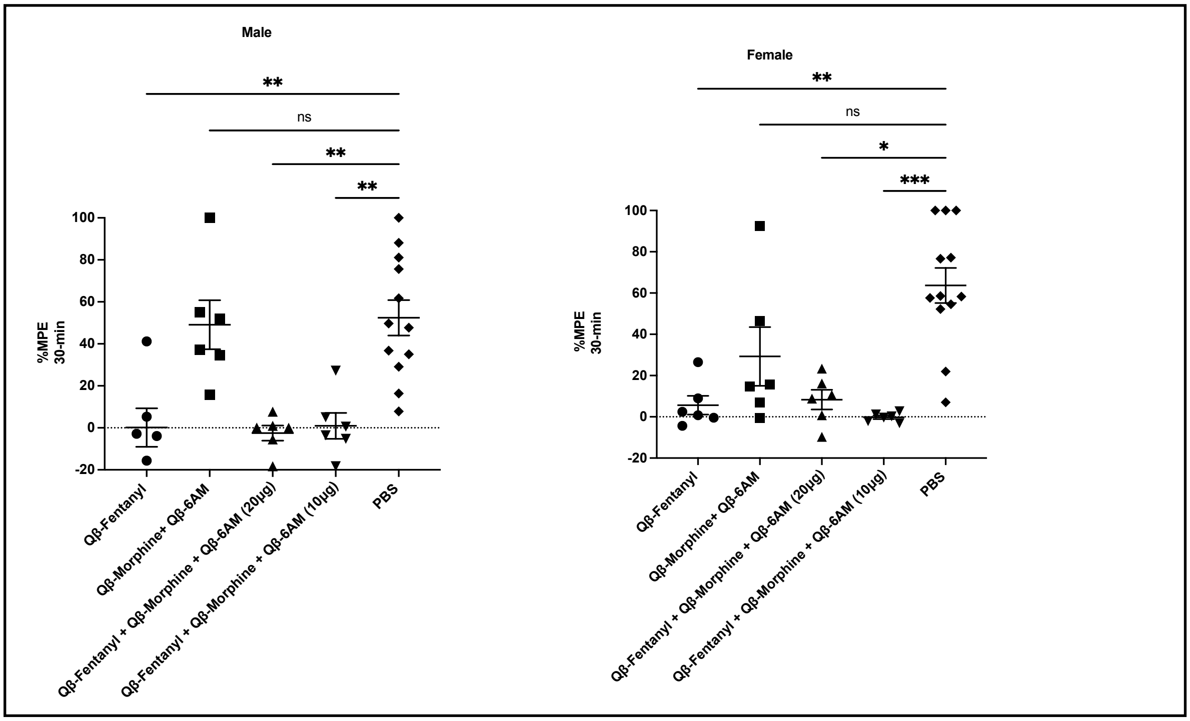


**Supplemental Figure 2. Both sexes display protection from fentanyl in a tail-flick challenge assay.** Tail- flick anti-nociceptive responses in vaccinated male (left) and female (right) mice (n=6, BALBc/j) challenged with s.c. fentanyl (0.0625 mg/kg) at three-weeks post-second immunization. Data shown as mean and SEM. % Maximum Possible Effect (%MPE=[(drug – basal response)/(20 sec – basal response time)] x 100%). *p<0.05, **p< 0.01,***p<0.001; Kruskal-Wallis test with Dunn's multiple comparison; GraphPad Prism.

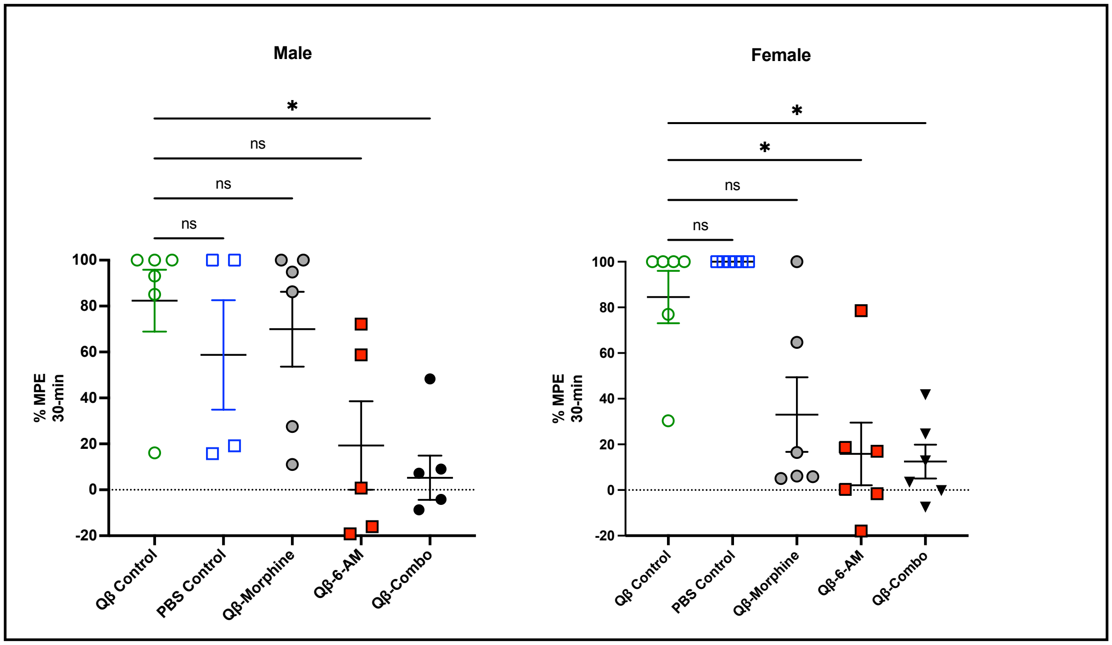


**Supplemental Figure 3. Both sexes display protection from heroin in a tail-flick challenge assay.** Tail-flick anti-nociceptive responses in vaccinated male (left) and female (right) mice (n=6, BALBc/j) challenged with s.c. heroin (0.5 mg/kg) at three-weeks post-second immunization. Qβ-Combo = Qβ-Morphine + Qβ-6-AM. Data shown as mean and SEM. % Maximum Possible Effect (%MPE=[(drug – basal response)/(20 sec – basal response time)] x 100%). $^*p<0.05$; Dunn's multiple comparison test; GraphPad Prism.
